# Phosphorylation of MdWRKY70L by MdMPK6/02G mediates reactive oxygen synthesis to regulate apple fruit senescence

**DOI:** 10.1101/2024.11.11.623016

**Authors:** Hui Wang, Shuhui Zhang, Yuchen Feng, Lulong Sun, Peng Yan, Yifeng Feng, Zhengyang Zhao

## Abstract

Apple (Malus domestica) is a globally significant crop and a vital dietary component worldwide. During ripening, apples exhibit a longitudinal gradient, ripening first at the stalk cavity and extending toward the calyx pit. Over-ripening in the stalk cavity leads to early senescence, characterized by peel browning, which diminishes fruit quality. This study examines the natural senescence process in 6-year-old ‘Ruixue’ apples by screening transcriptome data to uncover senescence-related genes and validate their molecular functions. Our analysis of antioxidant capacity and reactive oxygen species in different peel regions revealed that malondialdehyde, hydrogen peroxide, and superoxide anion levels increased with senescence, especially in the stalk cavity. Transcriptome clustering and enrichment analyses across developmental stages revealed *MdWRKY70L*, *MdSAG101*, and *MdZAT12* as key regulators of peel senescence. Further in *vitro* and in *vivo* studies demonstrated that MdWRKY70L is phosphorylated at Ser199 by MdMPK6/02G, enhancing its stability and promoting peel senescence. These findings offer insights for developing strategies to delay fruit senescence and improve postharvest quality control.

## INTRODUCTION

Apple (*Malus domestica* Borkh.) ranks among the top economically valuable crops globally. Its fruits are rich in essential vitamins, antioxidants, and cellulose, which are crucial components of the human diet and key to the global fruit trade (Zhao et al., 2020; Wang et al., 2023). However, the large-scale production and concentrated harvest times often result in delayed harvesting, pushing apples rapidly into natural senescence and triggering programmed cell death (Tian et al., 2013; Wang et al., 2023). Research on tomatoes has unveiled that fruit maturation follows a longitudinal gradient, beginning at the peduncle and advancing toward the stalk cavity, with coordinated genetic, hormonal, and metabolic changes along this axis (Shinozaki et al., 2018; Huang et al., 2022). Our previous investigation observed similar patterns in apples, where ripening initiates in the stalk cavity. By full maturity, the stalk cavity begins senescing, showing browning spots that spread outward, reducing the fruit’s visual appeal, market value, and shelf life (Wang et al., 2023). Despite these patterns being recognized, the molecular mechanisms underlying apple senescence remain poorly understood. Clarifying these regulatory mechanisms is crucial for advancing high-quality, efficient apple production.

Natural senescence marks the final stage of plant growth, driven by complex physiological and biochemical processes. It is primarily caused by an imbalance in reactive oxygen species (ROS) production and clearance within plant cells, leading to excessive ROS accumulation and subsequent oxidative damage (Mittler et al., 2022; Wang et al., 2024; Zhu et al., 2023). In *Arabidopsis*, ROS amassing promotes leaf senescence (Yang et al., 2018), and this process is affected by the gene *AtWRKY75*, which inhibits CAT2 degradation, thereby modulating ROS levels in *vivo* (Guo et al., 2017). In fruits such as tomato, kiwi, grape, peach, pear, loquat, and litchi, superoxide anion (O_2_^−^·) and hydrogen peroxide (H_2_O_2_) levels often rise 2- to 3-fold or more during ripening and senescence (Tian et al., 2013). ROS also function as signaling molecules, activating senescence-related genes like *SAGs*, *PAOs*, *CCGs*, *AAOs*, *LOXs*, *FADs*, *SODs*, *LEAs*, *PODs*, *PALs*, *CADs*, *PPOs*, and *LACs*. This gene activation results in senescence symptoms: chlorophyll degradation, decreased photosynthetic activity, yellowing leaves, dull fruit coloration, and the appearance of brown spots on the fruit peel (Zhang et al., 2018; Chen et al., 2023). Although senescence mechanisms have been widely studied in leaves, comparatively less research has focused on natural fruit senescence. Understanding this process more fully is essential for developing strategies to delay senescence and maintain fruit quality.

Plant maturation and senescence are regulated by numerous transcription factors that individually or cooperatively control specific downstream genes (Kuang et al., 2012; Shan et al., 2012; Xiao et al., 2013; Zhao et al., 2013). Recent studies have focused on WRKY transcription factors in plant senescence. For instance, in rice, OsWRKY42 suppresses *OsMT1d* expression, limiting ROS removal and accelerating leaf senescence (Han et al., 2014). Interaction of jasmonic acid (JA)-induced protein ESR with AtWRKY53 reduces its DNA-binding activity, leading to delayed senescence (Miao and Zentgraf, 2007). In *Arabidopsis*, various WRKY factors regulate senescence. For example, AtWRKY45 promotes natural senescence by modulating SAGs (Chen et al., 2017), AtWRKY57 inhibits senescence by repressing *SEN4/SAG12* (Jiang et al., 2014), and AtWRKY6 affects both senescence and pathogen defense through the senescence-induced receptor kinase pathway (Robatzek and Somssich, 2002). Our study uncovered *MdWRKY70L* as a key modulator of apple fruit senescence through transcriptomic analysis. Molecular tests, including transient injection, stable overexpression, and CRISPR/Cas9 knockout, demonstrated that *MdWRKY70L* promotes fruit senescence. These findings offer new insights into WRKY transcription factors’ roles in apple fruit senescence, opening pathways for future research and potential interventions to manage fruit senescence.

Beyond transcriptional regulation, the mitogen-activated protein kinase (MAPK) signaling cascade is critical for plant growth and development, with WRKY transcription factors as key downstream substrates (Sun and Zhang, 2022). For example, the overexpression of AtWRKY53 promotes senescence, and MEKK1 phosphorylates WRKY53, enhancing its DNA-binding ability. Moreover, WRKY53 can bind to its own promoter region, allowing it to be expressed not only during leaf senescence but throughout the plant senescence process (Miao et al., 2007). In our study, we also discovered that MdMPK6/02G interacts with MdWRKY70L, with phosphorylation modulating MdWRKY70L activity and enhancing its stability. This interaction affects ROS levels in fruits, ultimately regulating the fruit senescence process. These findings provide robust evidence for transcriptional and posttranslational regulation mechanisms related to fruit senescence and offer valuable insights for strategies aimed at maintaining postharvest fruit quality and extending storage time.

## RESULTS

### Ultrastructure and ROS dynamics during apple fruit senescence

Starch staining effectively indicates fruit ripening and senescence, showing that apple maturation follows a top-to-bottom gradient. Ripening begins in the stalk cavity, moves to the fruit surface, and finally reaches the calyx pit (Figure 1a). As maturation progresses, the stalk cavity initiates senescence, marked by increasing peel browning severity (Figure 1a, b). Chlorophyll content also declines gradually across fruit regions, with the steepest reduction in the stalk cavity (Figure S1a). Analyses of antioxidant and ROS dynamics revealed a progressive decline in total phenols, flavonoids, flavanols, and overall antioxidant capacity during senescence, especially in the stalk cavity (Figure S1b-e). Meanwhile, levels of malondialdehyde, H₂O₂, and O₂⁻· in the peel increased during fruit senescence, particularly in the stalk cavity (Figure 1c-e), where ROS-scavenging enzyme activity was notably reduced (Figure S1f-h). Ultrastructure examination showed that, in unripe fruits, peel cells in the stalk cavity, surface, and calyx pit exhibited a honeycomb-like structure with smooth, intact cells (Figure 1f). However, with senescence onset, especially in the stalk cavity and surface, cells showed deformation, cell wall thickening, subepidermal cells sinking, and general tissue disorganization, losing the honeycomb pattern (Figure 1g). Moreover, starch particles were nearly absent in the cells at the browning sites, whereas the number of osmiophilic droplets increased, and chloroplast degradation was evident. By contrast, cells in the calyx pit, which exhibited no browning, retained visible starch particles, had fewer osmiophilic droplets, and displayed intact chloroplast structure, with minimal degradation of the thylakoid grana (Figure 1h). These findings indicated that apple fruit follows a pattern of longitudinal gradient maturation, beginning at the stalk cavity and extending toward the calyx pit. At full maturity, the fruit enters the senescence stage, starting from the stalk cavity. The decrease in antioxidant capacity, cell damage, and a significant increase in ROS are the primary factors contributing to fruit senescence.

**Figure 1.**
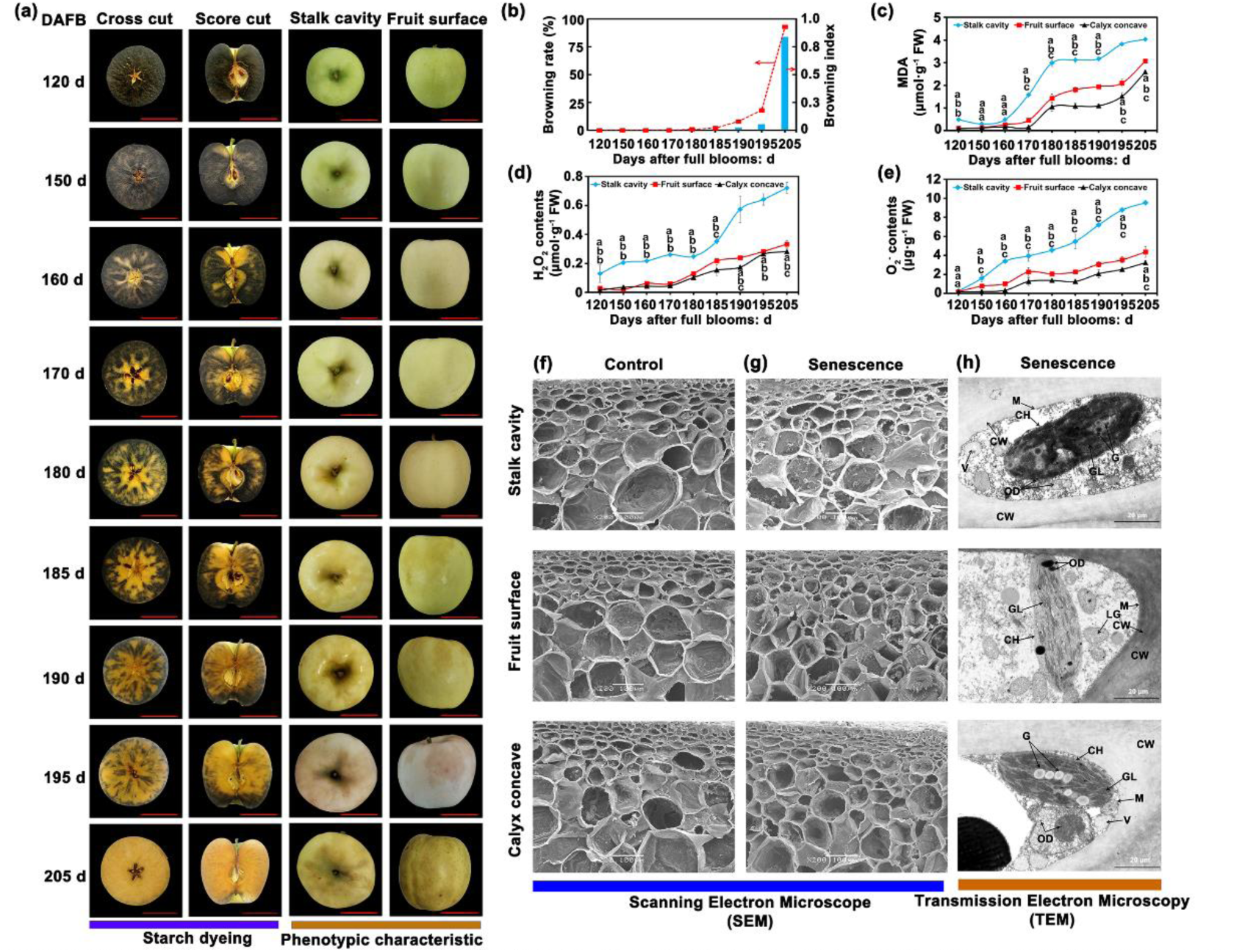
Changes in senescence characterization, ROS system, and ultrastructure in different parts of apple fruits during senescence. **(a)** Starch dyeing and phenotypic analysis of apples at different developmental stages. Digital images were extracted for comparison. Bars = 4 cm. **(b)** Analysis of senescence browning index and rate of apples at different stages (n = 300 fruits). **(c-e)** contents of MDA, O_2_^-^·, and H_2_O_2_. The X-axis indicates sampling time. Data presented as mean ± SD with nine fruits per measurement. **(f-h)** Ultrastructure of the non-senescent and senescent cell wall (CW), vacuole (V), chloroplast (CH), lipid globules (LG), mitochondria (M), osmiophilic droplets (OD), grain (G), grana lamella (GL), intercellular space (ICS), and vesica (Ve).

### *MdWRKY70L* as a key regulator of apple fruit senescence

Research has shown that WRKY transcription factor family genes are integral to regulating senescence in crops (Zhou et al., 2011). To investigate WRKY genes linked to apple peel senescence, we performed clustering and enrichment analysis on WRKY family genes using transcriptome data across different developmental stages. In total, 31 WRKY family genes exhibited differential expression. Among these, the expression levels of *MdWRKY1* (MD09G0105800), *MdWRKY3* (MD13G0059600), *MdWRKY31* (MD03G0162000), *MdWRKY24* (MD03G0048200), *MdWRKY48* (MD13G0134000), *MdWRKY65* (MD05G0248800), *MdWRKY69* (MD09G0202900), *MdWRKY70L* (MD01G0136400), *MdWRKY72A* (MD13G0068300), *MdWRKY75* (MD13G0108800), and *MdWRKY76* (MD15G0034900) exhibited a gradually increasing expression as fruit senescence progressed (Figure 2a). Further reverse transcription quantitative polymerase chain reaction (RT-qPCR) analysis confirmed that MdWRKY70L exhibited the highest differential expression among these genes (Figure 2b), suggesting that the elevated *MdWRKY70L* expression is strongly associated with fruit senescence.

**Figure 2.**
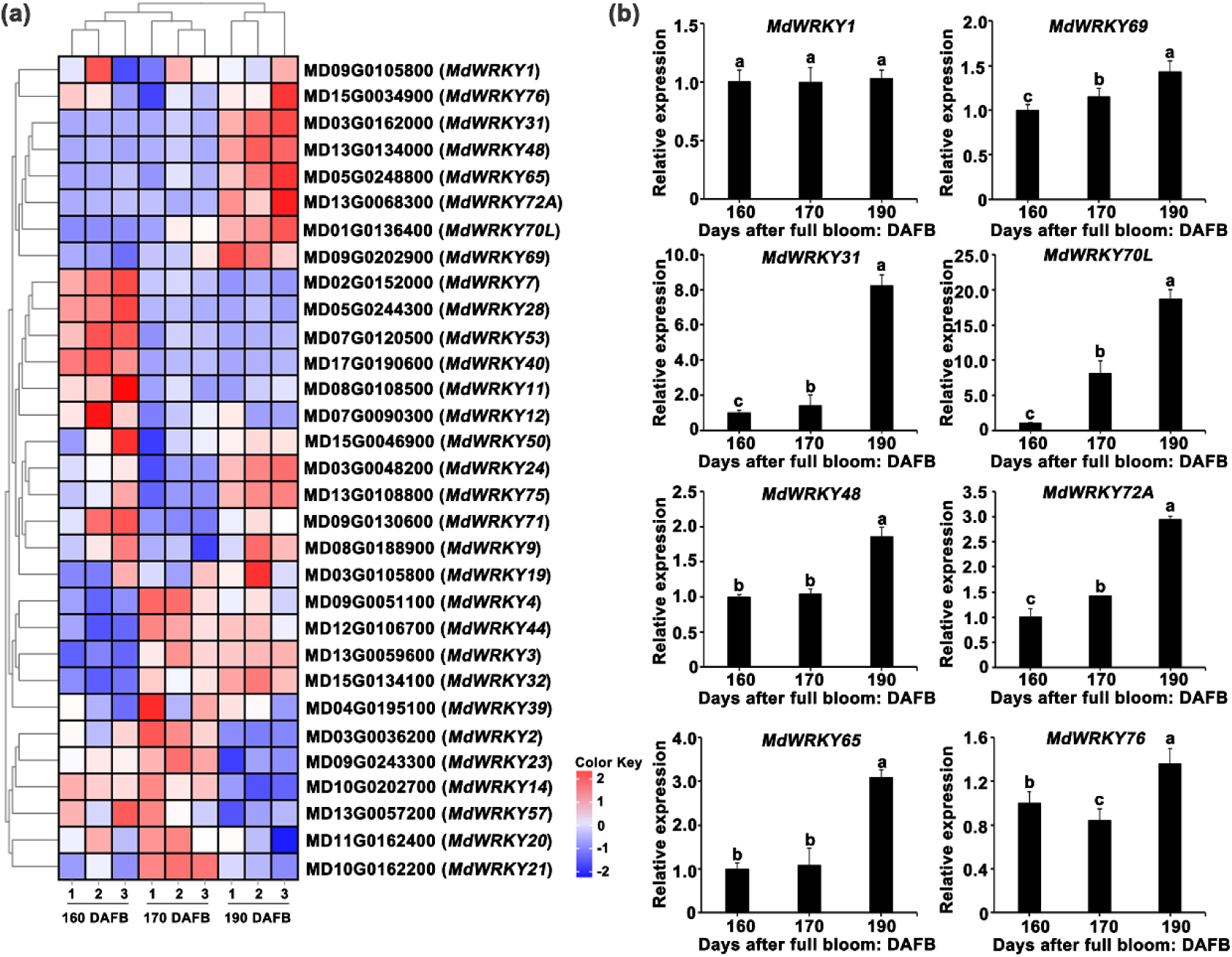
Screening of *MdWRKY70L* transcription factors. **(a)** Expression profiles of *MdWRKY* family genes. **(b)** Differentially expressed genes *MdWRKY1*/*3*/*31*/*24*/*48*/*65*/*69*/*70L*/*72A*/76 were identified. Data shown are mean ± standard error with different letters denoting *P* < 0.05 (Student’s *t* test).

### *MdWRKY70L* promotes senescence in apple and ‘orin’ calli

To assess the role of *MdWRKY70L* in apple fruit senescence, we constructed *MdWRKY70L* overexpression vectors (pCAMBIA2300-*MdWRKY70L*) and silencing vectors (pTRV2-*MdWRKY70L*), which were transiently transformed into apple peel tissue by using *Agrobacterium tumefaciens* as a mediator. Overexpression of *MdWRKY70L* significantly increased *MdWRKY70L* expression and accelerated senescence on the peel surface compared with the empty vector control. By contrast, no senescence phenotype was observed following *MdWRKY70L* silencing (Figure 3a, b). ROS assays showed that in peel tissues overexpressing *MdWRKY70L*, the contents of O₂⁻· and H₂O₂ increased by 24.5% and 32.4%, respectively, compared with the PC23300-GFP control. By contrast, in peel tissues with *MdWRKY70L* silencing, O₂⁻· and H₂O₂ levels were significantly reduced by 21.7% and 32.6%, respectively (Figure 3c, d). We further confirmed these results in ‘Orin’ calli with stable overexpression and knockout of *MdWRKY70L* (Figure 3e-h). In the overexpressing calli, *MdWRKY70L* expression was significantly increased (Figure 3g), and O₂⁻· and H₂O₂ levels rose by 0.7- and 2.6-fold, respectively, compared to wild-type (WT) calli, showing severe senescence (Figure 3i-k). In knockout calli, *MdWRKY70L* expression was nearly undetectable, with O₂⁻· and H₂O₂ contents reduced by 61.8% and 58.8%, respectively, resulting in a youthful appearance with no senescence signs (Figure 3i-k). These findings demonstrate that *MdWRKY70L* is essential for driving the senescence process in both apple fruit and ‘Orin’ calli.

**Figure 3.**
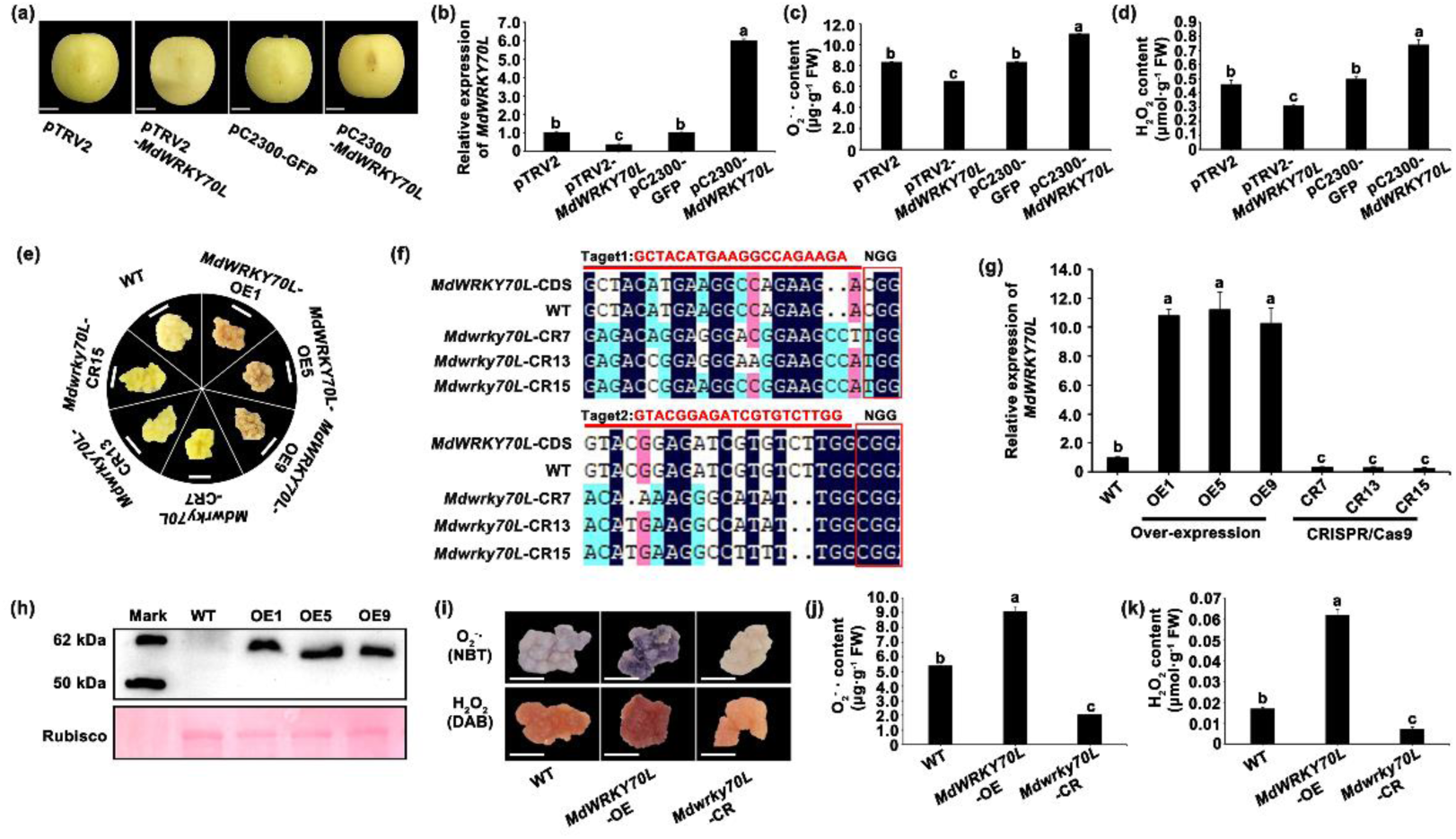
Functional validation of *MdWRKY70L* transcription factor. **(a)** Effects of *MdWRKY70L* on apple phenotypes, with digital images for comparison. Bars = 2 cm. **(b)** *MdWRKY70L* expression and **(c)** O_2_^−^·and **(d)** H_2_O_2_ contents in apple peels post-transient *MdWRKY70L* integration. Dats shown are mean ± standard error with different letters denoting *P* < 0.05 (Student’s *t* test). **(e)** Phenotype of *MdWRKY70L* overexpressing ‘Orin’ calli (OE-*MdWRKY70L*-1/5/9) and CRISPR/Cas9 knockdown calli (CR-*MdWRKY70L*-7/13/15). **(f)** Sequence verification of transgenic knockout materials. **(g-h)** *MdWRKY70L* RNA **(g)** and protein **(h)** levels in stable overexpressing lines. **(i)** NBT and DAB staining results of ‘Orin’ calli with *MdWRKY70L* stable overexpression and CRISPR/Cas9-mediated knockout. **(j)** O_2_^-^· content. **(k)** H_2_O_2_ content. Data shown are mean ± standard error with different letters denoting *P* < 0.05 (Student’s *t* test).

### *MdWRKY70L* regulates senescence-related genes in apple fruits

To uncover the regulatory mechanism of *MdWRKY70L* in fruit senescence, we analyzed DEGs from transcriptome data. This analysis revealed that SAGs, programmed cell death family genes, and genes involved in salicylic acid, ethylene, abscisic acid, JA, and ROS biosynthesis were significantly enriched (Figure 4a). RT-qPCR analysis further revealed that the levels of *MdSAG101* (MD09G0034000), *MdEDS1* (MD14G0164000), *MdCBP60F* (MD12G0174000), *MdCYP76B6* (MD13G0103200), *MdACO1* (MD17G0093500), *MdACS1* (MD14G0097100), *MdAAO1* (MD11G0144200), *MdLOX1.5* (MD04G0166700), and *MdZAT12* (MD07G0159300) increased as peel senescence progressed (Figure S2). Correlation analyses showed significant positive correlations between *MdWRKY70L* and both *MdSAG101* and *MdZAT12*, with correlation coefficients of 0.99 and 0.97, respectively (Figure 4b), suggesting that *MdWRKY70L* accelerates peel senescence by regulating *MdSAG101* and *MdZAT12*, as their expression was augmented after transient *MdWRKY70L* injection and in stable *MdWRKY70L* transgenic ‘Orin’ calli. We found that the overexpression of *MdWRKY70L* promoted their expression, whereas silencing or knocking out *MdWRKY70L* significantly inhibited their expression. Notably, *MdSAG101* and *MdZAT12* exhibited the largest difference in variation (Figure S3a, b). To further validate the roles of *MdSAG101* and *MdZAT12* in fruit senescence, we transformed the pTRV2-*MdSAG101*/pTRV2-*MdZAT12* gene silencing vectors and the pCAMBIA2300-*MdSAG101*/pCAMBIA2300-*MdZAT12* overexpression vectors into apple peel tissue, using *Agrobacterium* as the mediator. Compared to controls, overexpressing *MdSAG101* and *MdZAT12* significantly augmented O₂⁻· and H₂O₂ levels in the peel, whereas silencing these genes reduced O₂⁻· and H₂O₂ contents (Figure 4c-g). In ‘Orin’ calli with stable overexpression of *MdSAG101* and *MdZAT12*, O₂⁻· and H₂O₂ levels were also significantly higher than in the WT (Figure 4h-i). These observations suggested that *MdWRKY70L* regulates *MdSAG101* and *MdZAT12* expression, thereby mediating ROS production in the peel and accelerating the fruit senescence process.

**Figure 4.**
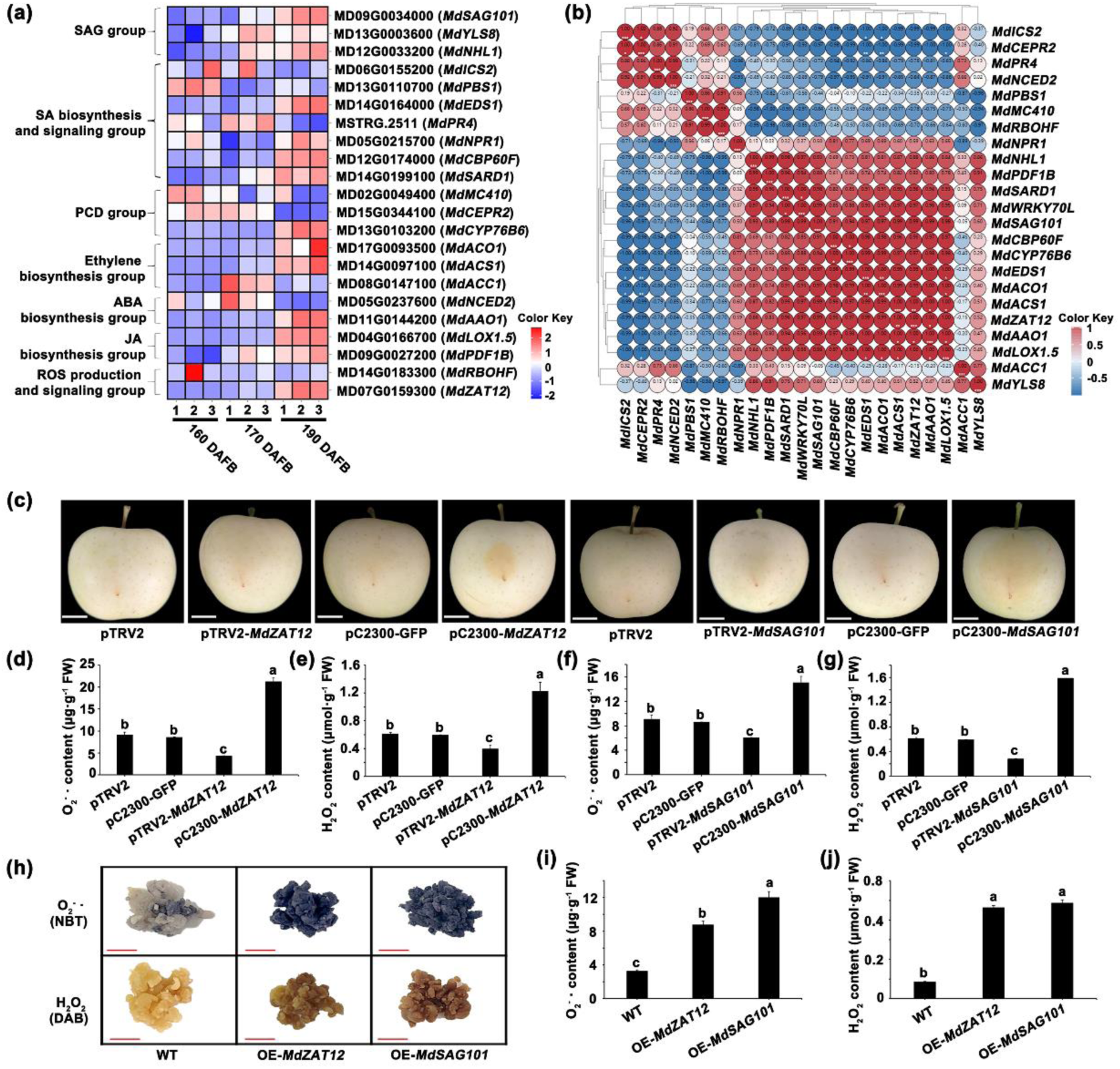
Screening and functional verification of senescence-related genes. **(a)** Expression profiles of senescence-related genes in apples at different developmental stages. **(b)** Correlation of *MdWRKY70L* with senescence-related genes. **(c)** Phenotypes of apples after transient *MdSAG101* and *MdZAT12* infection. Digital images were isolated for comparison. Bars = 2 cm. **(d–g)** O_2_ ^-^·and H_2_O_2_ contents in apple after instant infection with *MdSAG101* and *MdZAT12*. Data shown are mean ± standard error with different letters denoting *P* < 0.05 (Student’s *t* test). **(h)** Phenotype of ‘Orin’ calli stably overexpressing *MdSAG101* and *MdZAT12* after staining with NBT and DAB. **(i)** O_2_^-^·content. **(j)** H_2_O_2_ content.

### *MdWRKY70L* positively regulates *MdSAG101*/*MdZAT12* expression to promote apple fruit senescence

To investigate how *MdWRKY70L* promotes senescence, we analyzed the promoters of *MdSAG101* and *MdZAT12* and identified W-box motifs (WRKY binding sites, TTGACC/CTGACT). Electrophoretic mobility shift assay (EMSA) uncovered that *MdWRKY70L* binds specifically to the W-box motif on the *MdSAG101* and *MdZAT12* promoters (Figure 5a, b). Additionally, luciferase (LUC) assays demonstrated *MdWRKY70L* binding to these promoters activates their expression in plant cells (Figure 5c-f).

**Figure 5.**
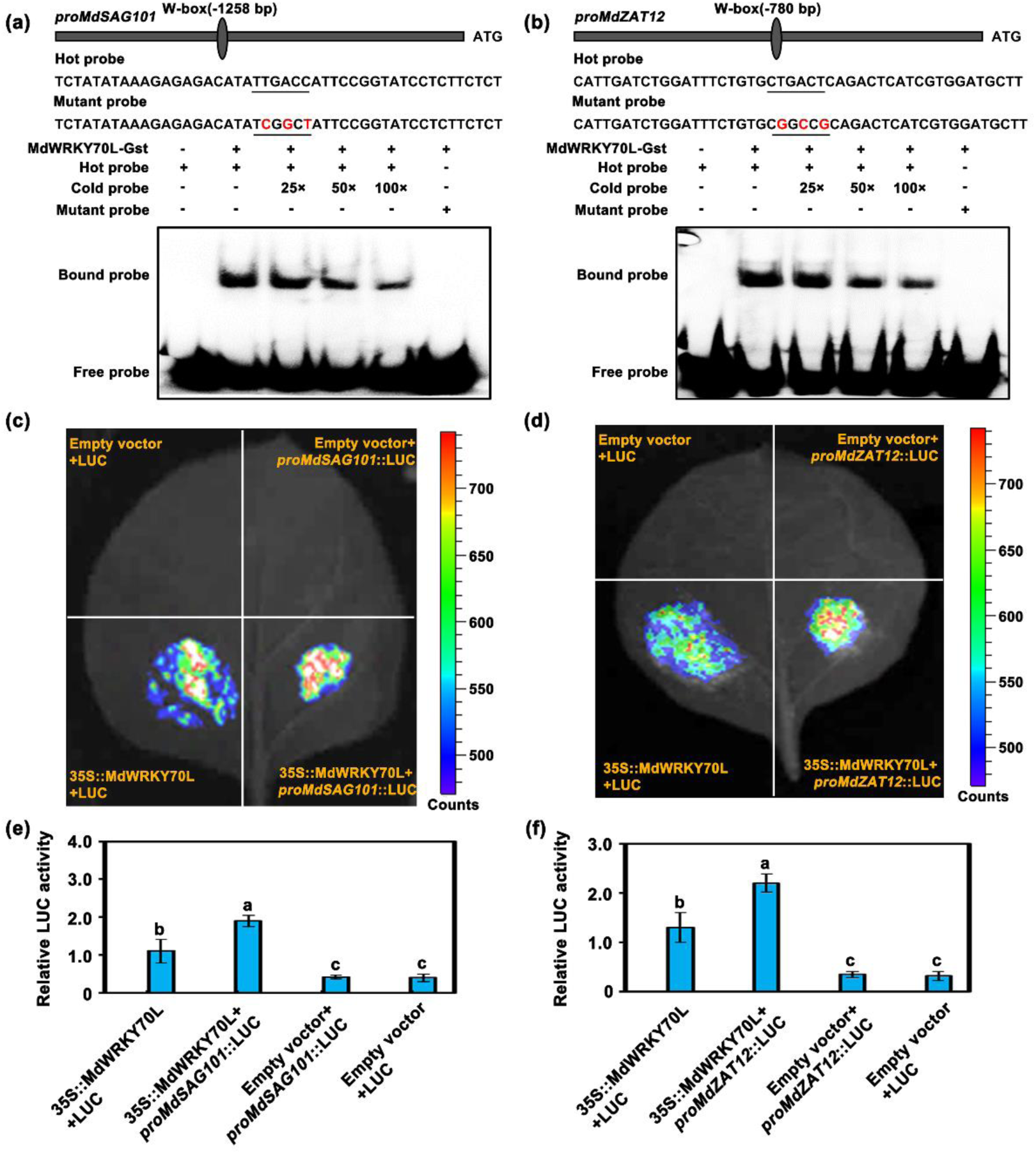
MdWRKY70L binding to *MdSAG101*/*MdZAT12’s* promoter. **(a, b)** Electrophoretic mobility shift assay using biotin-labeled and unlabeled promoter probes specific to *MdSAG101*/*MdZAT12*’s W-box motifs and a mutated probe indicating MdWRKY70*L’s* binding to *MdSAG101*/*MdZAT12* promoter. Cold probes were provided incrementally (25×, 50×, and 100×). The “+” and “-” symbols denote the inclusion and exclusion of each probe or protein, respectively. **(c-f)** Luciferase analysis uncovering *in vivo* MdWRKY70L binding to the *MdSAG101*/*MdZAT12* promoter in agroinfiltrated *Nicotiana benthamiana* leaves on day 3. Data shown are mean ± standard error with different letters denoting *P* < 0.05 (Student’s *t* test).

To explore if *MdWRKY70L* promotes fruit senescence by modulating *MdSAG101* and *MdZAT12*, we separately transformed *MdSAG101* and *MdZAT12* into *MdWRKY70L* overexpression and knockout ‘Orin’ calli. The results showed that in *MdWRKY70L* knockout calli, the overexpression of *MdSAG101* and *MdZAT12* induced senescence phenotypes, and the O₂⁻· and H₂O₂ contents significantly increased compared with the WT (Figure S4a-c). By contrast, in *MdWRKY70L* overexpression calli, the stable transformation of *MdSAG101* and *MdZAT12* intensified senescence phenotypes, with markedly higher O₂⁻· and H₂O₂ levels than the WT (Figure S4a-c). These observations demonstrated that *MdWRKY70L* acts in conjunction with *MdSAG101* and *MdZAT12* both in *vivo* and in *vitro* to accelerate apple fruit senescence. **MdMPK6/02G interacts with MdWRKY70L and enhances its protein stability** Protein modification, such as phosphorylation, is essential in regulating protein functions, with WRKY transcription factors often undergoing phosphorylation to facilitate plant growth and development. Using LC-MS/MS analysis on proteins extracted from *MdWRKY70L*-GFP transgenic ‘’Orin’ calli, we unveiled phosphorylated peptides in *MdWRKY70L*-GFP samples, confirming that MdWRKY70L undergoes phosphorylation (Figure S5a-b, Table S1). To further explore this, we performed yeast two-hybrid (Y2H) screening and observed associations between MdMPK6/02G and MdWRKY70L. Specifically, yeast cells co-expressing *MdWRKY70L*-PGAD and *MdMPK6/02G*-PGBK exhibited regular growth on the selective medium (−T/−L/−H/−A) and blue coloration, indicating interaction. Additional pull-down, bimolecular fluorescence complementation (BiFC), and luciferase complementation imaging (LCI) assays further validated this protein-protein interaction (Figure 6a-d). Subsequently, *MdMPK6/02G*-flag and *MdMPK6/02G*-TRV2 vectors were transformed into ‘’Orin’ calli, total protein was extracted, and purified MdWRKY70L-His protein was co-incubated at 22°C for 0, 1, and 3 h. Overexpression of *MdMPK6/02G* in the calli enhanced the stability of MdWRKY70L-GST protein over time compared with *MdMPK6/02G*-TRV2 and WT calli (Figure S6). These results indicated that MdMPK6/02G interacts with MdWRKY70L and enhances its protein stability.

**Figure 6.**
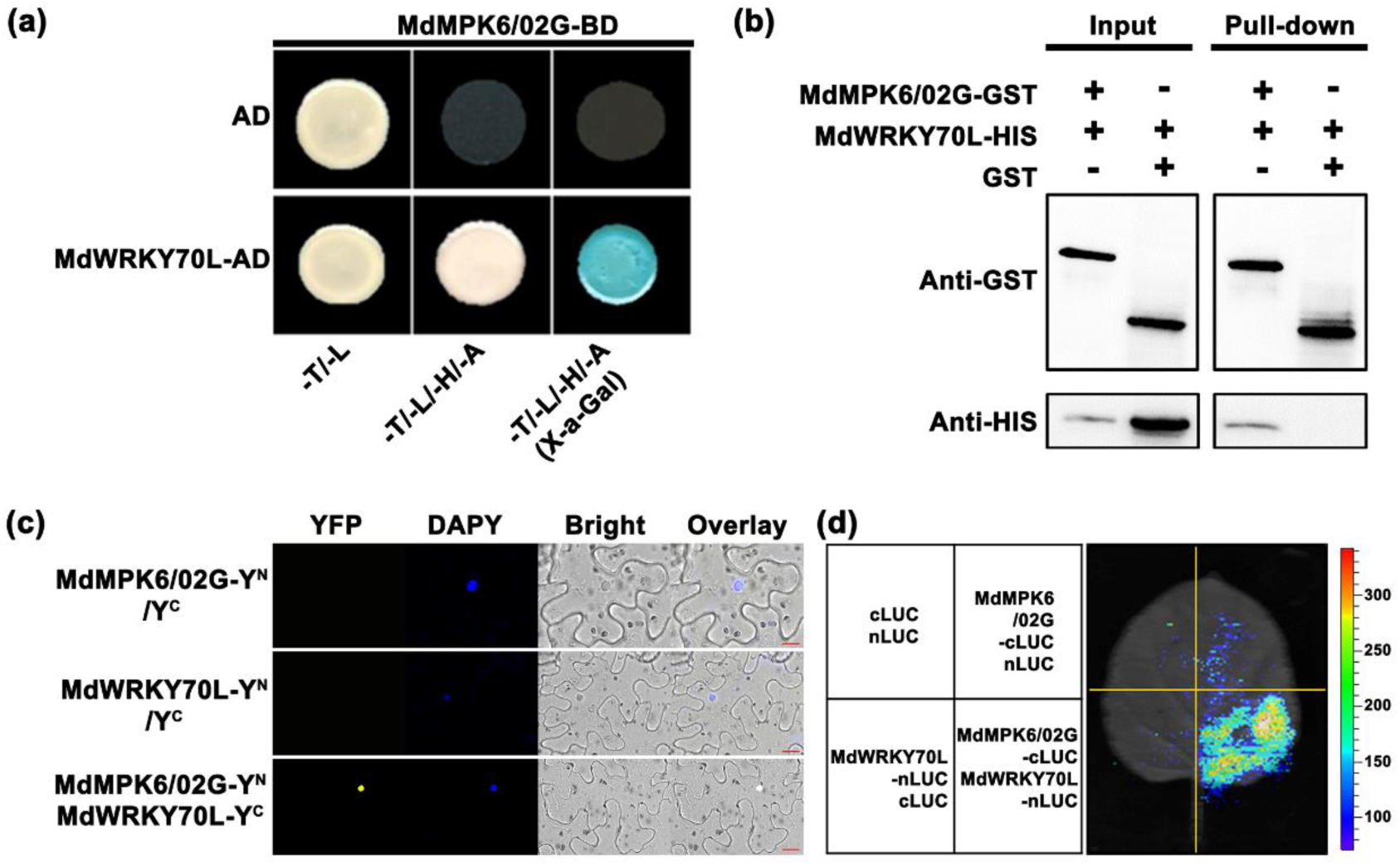
*In vivo and in vitro* interactions between MdMPK6/02G with MdWRKY70L enhances MdWRKY70L stability. The interaction was confirmed using the yeast two-hybrid assay **(a)**, pull-down assay **(b)**, bimolecular fluorescence complementation assay **(c)**, and luciferase complementation imaging **(d)**.

### MdMPK6/02G promotes fruit senescence by phosphorylating MdWRKY70L at **Ser199**

We identified phosphorylation at the Ser199 site of MdWRKY70L through immunoprecipitation and mass spectrometry (IP/MS) (Table S1). To verify the role of MdMPK6/02G in phosphorylating MdWRKY70L, we obtained active CAMdMPK6/02G-GST protein and a point mutant version of MdWRKY70L with a Ser199 mutation (MdWRKY70L*^s199^*-GST) for in *vitro* analysis. In an in *vitro* phosphorylation experiment using kinase buffer, we found that CAMdMPK6/02G could phosphorylate MdWRKY70L but was unable to phosphorylate MdWRKY70L^s199^, indicating that CAMdMPK6/02G regulates MdWRKY70L activity by phosphorylating it at the Ser199 site (Figure 7a).

**Figure 7.**
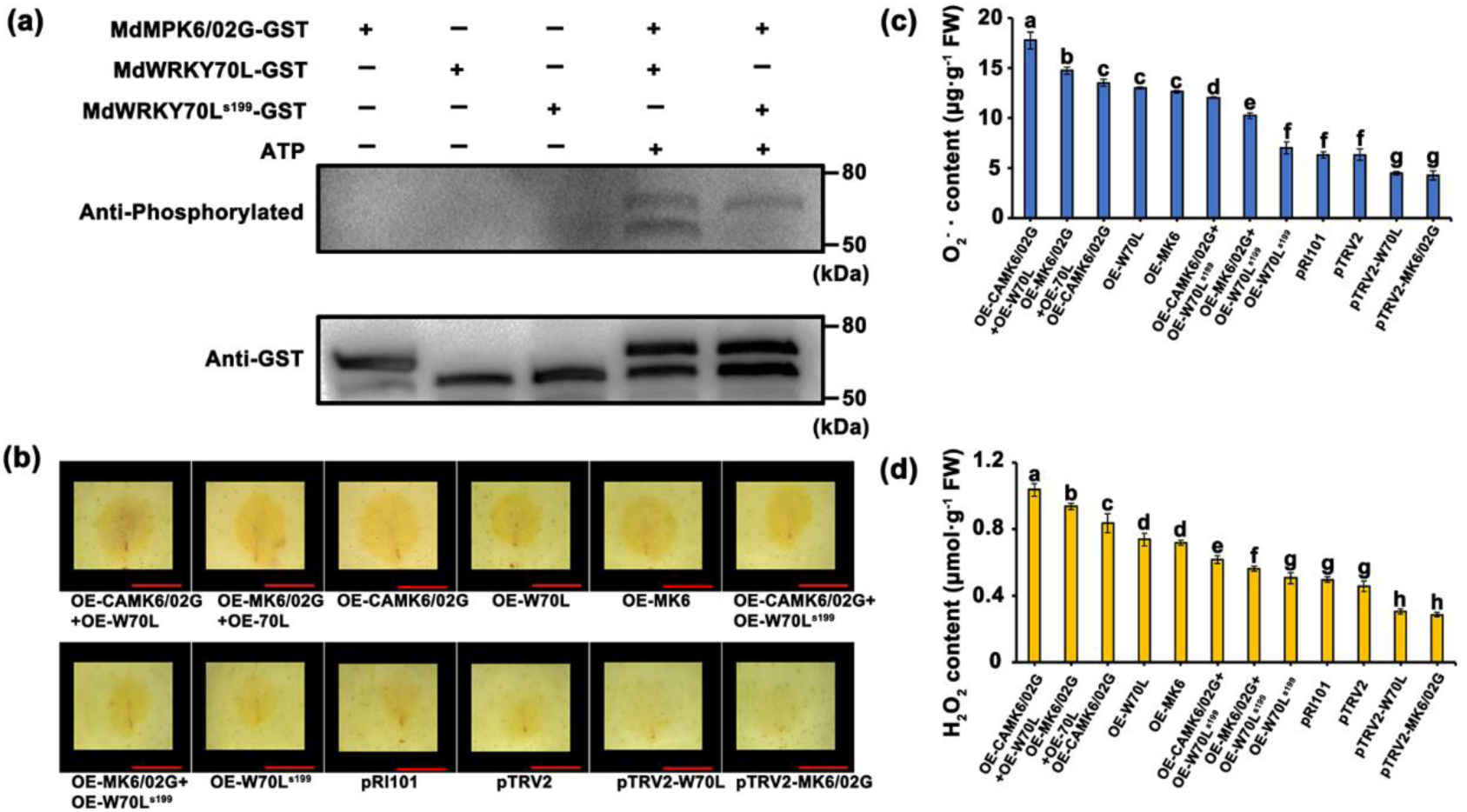
MdWRKY70L phosphorylation at Ser199 by MdMPK6/02G promotes fruit senescence. **(a)** *In vitro* phosphorylation modification. **(b)** Transient transgene integration verification for fruit senescence induced by phosphorylation of MdWRKY70L at Ser199 by MdMPK6/02G. Scale bar = 2 cm. Apple images were digitally processed for comparison. **(c, d)** O_2_^−^· and H_2_O_2_ contents in apples after instant infection with *MdMPK6/02G and MdWRKY71.* Data shown are mean ± standard error with different letters denoting *P* < 0.05 (Student’s *t* test).

Next, we conducted transient transformation experiments, overexpressing *MdWRKY70L*-GFP, *MdWRKY70L^s199^*-GFP, *CAMdMPK6/02G*-flag, *CAMdMPK6/02G*-flag + *MdWRKY70L*-GFP, and *CAMdMPK6/02G*-flag + *MdWRKY70L^s199^*-GFP in the apple peels. The results showed that co-transfection of *MdMPK6/02G*-flag with *MdWRKY70L*-GFP in the apple peel led to severe senescence phenotypes (Figure 7b, Figure S7), with a significant increase in O₂⁻· and H₂O₂ levels (Figure 7c, d). However, when the Ser199 site of *MdWRKY70L* was mutated, co-transfection of *MdMPK6/02G*-flag with *MdWRKY70L^s199^*-GFP alleviated the symptoms of peel senescence (Figures 7b and S7). Moreover, the O₂⁻· and H₂O₂ levels were significantly decreased (Figure 7c, d). These effects were even more pronounced when *CAMdMPK6/02G*-flag was expressed. In conclusion, MdMPK6/02G phosphorylates MdWRKY70L at the Ser199 site, thereby promoting senescence in apple peel.

## DISCUSSION

Fruit growth and development can proceed through five stages: cell differentiation, cell expansion, fruit development, ripening, and senescence, with natural senescence being the final stage. This stage is crucial as it directly affects fruit quality, market value, and shelf life (Giovannoni, 2001). Fruit senescence is a complex, highly regulated physiological and biochemical process, tightly regulated and influenced by ROS accumulation (Buchanan-Wollaston et al., 2005; Zhu et al., 2018). As senescence progresses, physiological functions decline, cell damage occurs, and pulp browning and decreased resistance to pathogens make the fruits more susceptible to spoilage, thus significantly shortening postharvest life and preservation time (Tian et al., 2013; Wang et al., 2005; Wang et al., 2023). In our study, ROS levels in various senescent fruit regions showed a progressive increase in O₂⁻· and H₂O₂. Meanwhile, antioxidant enzyme activities and antioxidant compound levels declined, confirming that ROS accumulation is a major factor mediating fruit senescence. While these findings align with earlier research, most studies on plant senescence mechanisms have focused on leaves. Further explorations are needed to clarify the specific dynamics of ROS changes during fruit senescence.

WRKY transcription factors are essential in plants, where they regulate gene expression by binding to W-box elements in promoter regions. These transcription factors function as either activators or repressors, influencing a range of processes, such as growth, responses to biotic and abiotic stresses, and hormone signaling (Wang et al., 2023). In fruit senescence, WRKY transcription factors often exhibit significant expression changes. For instance, in banana fruits, *MaWRKY31* activates the promoter activity of ethylene synthesis genes *MaACS1* and *MaACO1*, which may enhance ethylene synthesis and accelerate fruit senescence (Xiao et al., 2013). Similarly, in tomato fruits, several *SlWRKY* genes are upregulated during fruit maturation and contribute to post-ripening regulation by controlling ethylene synthesis, pigment accumulation, fruit softening, and other related processes (Huang et al., 2022). Beyond ethylene synthesis, WRKY factors can directly target senescence-associated genes, such as MaWRKY31’s activation of *MaSAG1* in banana (Xiao et al., 2013), and modulate ROS levels, thereby mediating plant senescence (Chen et al., 2017). In this study, MdWRKY70L was observed to bind to W-box motifs in the promoters of *MdSAG101* and *MdZAT12*, actively regulating their expression, thereby affecting ROS levels and promoting fruit senescence. These results deepen our understanding of the transcriptional regulation pathways that control fruit senescence.

WRKY transcription factor activity is primarily modulated through MAPK-mediated phosphorylation. In *Arabidopsis*, AtWRKY33 phosphorylation by MPK3/MPK6 regulates plant antitoxin biosynthesis (Mao et al., 2011). Similarly, the absence of an MPK3/MPK6 phosphorylation site affects WRKY34 function *in vivo* (Guan et al., 2014). *OsWRKY53* negatively modulates MPK3/MPK6 to activate early plant defense responses (Hu et al., 2015) and interacts with the OsMAPKK4*-*OsMAPK6 cascade to influence brassinolide signaling (Tian et al., 2017). In addition, MPK1 phosphorylation of WRKY53 in *Arabidopsis* enhances its DNA-binding ability, accelerating the leaf senescence process (Li et al., 2020). Our study unveiled that MdMPK6/02G phosphorylates MdWRKY70L at Ser199 to enhance its stability. This modification further promotes the regulation of the downstream senescence-related genes *MdSAG101* and *MdZAT12*, leading to increased ROS accumulation and ultimately causing fruit senescence and browning (Figure 8). These findings offer promising potential for molecular-assisted breeding to delay fruit senescence and preserve fruit quality.

**Figure 8.**
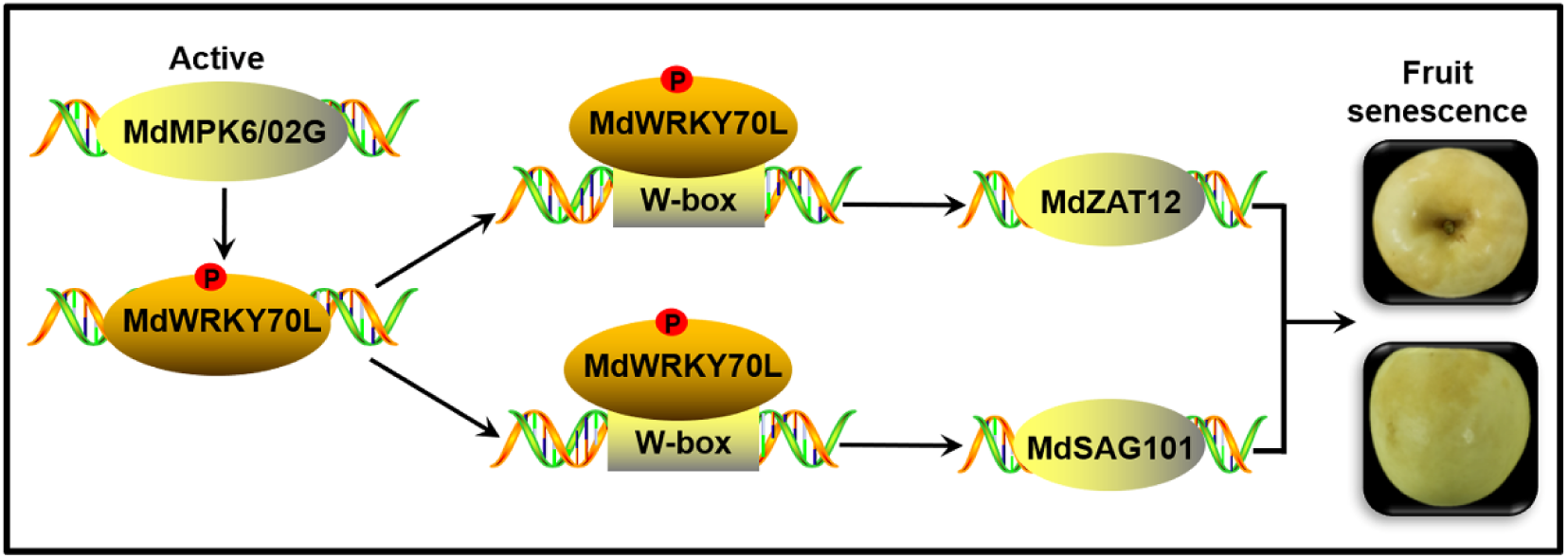
Proposed model for apple fruit senescence regulated by MdMPK6/02G-mediated MdWRKY70L phosphorylation and ROS synthesis. *MdWRKY70L* TF is a key candidate gene that regulates apple fruit senescence. MdWRKY70L could interact with *MdSAG101*/*MdZAT12* both in *vivo* and in *vitro*, thereby mediating ROS production in peel and accelerating the fruit senescence process. In addition, MdMPK6/02G phosphorylates and stabilizes MdWRKY70L, further promoting senescence phenotype in apples.

## MATERIALS AND METHODS

### Plant materials and treatments

This study used six-year-old ‘Ruixue’ apple trees and ‘Orin’ calli as test materials. The experiment took place from June to November 2022 at the Bai Shui Apple Experimental Station (35°02’N, 109°06’E, 908 m altitude) of Northwest A&F University. The site, located in a moderate monsoon climate with continental features, experiences a mean annual rainfall and daily mean temperature of 578 mm and 11.4°C, respectively.

To prepare samples, fruits were routinely bagged 55 days post after full bloom (DAFB; June 15). Sampling began at 160 DAFB and continued at 10-day intervals across four sampling points. At each time, 30 fruits of similar size, maturity, and without mechanical damage were selected from each group. The peel was carefully removed using a sterile scalpel, rapidly frozen in liquid nitrogen, and kept at −80°C for later experiments.

For genetic transformation, ‘Orin’ calli were cultured on MS medium supplemented with 1.5 mg L^-1^ 2,4-dichlorophenoxyacetic acid (2,4-D) and 0.4 mg L^-1^ 6-benzylaminopurine in the dark at 25°C and refreshed every 20 days. Meanwhile, *Nicotiana benthamiana* was cultivated under a 16 h/8 h light/dark cycle at 25°C and 70% ± 5% relative humidity in a light incubator.

### Browning rate and index

The browning rate and index were assessed using a slightly modified method from Wang et al. (2023). The browning rate is the percentage of browned fruits in a sample of 300 randomly selected fruits. Browning severity (S) was rated on a 0−3 scale with 0, 1, 2, and 3 for no, mild (<1/3 of the fruit’s surface), moderate (between 1/3 and 2/3), and severe (> 2/3) browning, respectively. The browning index was determined as ∑ [(browning scale) × (fruit count at that scale)] / (3 × total fruit count) × 100.

### Antioxidant capacity

Total antioxidant activity and components were measured using a modified method based on Wang et al. (2023). 0.5 g of fresh peel samples were prepared as a fine powder and mixed with 1.5 mL of a 7:3 (v/v) ethanol-acetone solution. The mixture was kept at 37°C for 1 h and spun for 10 min at 13,000 rpm and 20°C. The resulting supernatant was instantly placed at −20°C for subsequent antioxidant capacity analysis. The experiments were conducted in triplicate, with three biological replicates for accuracy.

### Histological staining for ROS detection

O₂⁻· and H₂O₂ levels were detected as previously described, with slight modifications (Zhang et al., 2024). Briefly, ‘Orin’ calli were immersed in 1 mg·mL⁻¹ DAB staining solution for H₂O₂ or 1 mg·mL⁻¹ NBT staining solution (for O₂⁻·). The calli were incubated under shaking (20 rpm) at 25°C overnight in the dark. After incubation, the calli were preserved in a solution of ethanol and glycerol (4:1) until imaging was performed.

### Microstructure of the peel cells

For microstructural analysis, peel tissues (1.0 × 2.0 × 5.0 mm) were cut with a scalpel and rapidly fixed in 4% glutaraldehyde (v:v). The samples were vacuumed to ensure complete immersion in the fixative and left overnight. Following this, the samples were rinsed with phosphate buffer (PBS; 0.1 mol·L^−1^, pH 6.8) and fixed for 2 h in 1% osmium tetroxide. After five 5-min PBS washes, the samples were dehydrated using a graded ethanol series (30%, 50%, 70%, 80%, and 90%), with each concentration applied for 10 min, followed by three 10-min washes with 100% ethanol and embedding overnight in epoxy propane and SPI-81 medium. Ultrathin sections (90 nm) were cut with a Leica EM UC7 ultramicrotome (Leica, Germany), stained with uranyl acetate and lead citrate, and examined under a Hitachi HT7700 microscope (Hitachi, Japan) (Wang et al., 2023).

### RNA isolation and quantification

RNA was isolated with TRIzol (Invitrogen, Carlsbad, CA, USA), and its integrity was evaluated using Agilent 2100 Bioanalyzers (Agilent Technologies, Palo Alto, CA, USA) and agarose gel electrophoresis. Gene levels were analyzed by RT-qPCR with three biological replicates using SYBR Green Master Mix (SYBR Premix EX TaqTM, Dalian, China) on an ABI7500 RT-qPCR system (ABI, MA, USA) (Wang et al., 2023). Supplemental Table S2 lists all primer information.

### *MdWRKY70L*, *MdSAG101*, *MdZAT12*, and *MdMPK6/02G* overexpression or silencing in fruits

The transient overexpression vectors *MdWRKY70L*-pCAMBIA2300, *MdSAG101*-pCAMBIA2300, *MdZAT12*-pCAMBIA2300, and *MdMPK6/02G*-pCAMBIA1300 were constructed by cloning corresponding coding sequences (CDS) into either pCAMBIA2300 or pCAMBIA1300 vectors while silencing vectors *MdWRKY70L*-pTRV2, *MdSAG101*-pTRV2, *MdZAT12*-pTRV2, and *MdMPK6/02G*-pTRV2 were obtained inserting fragments specific to *MdWRKY70L*, *MdSAG101*, *MdZAT12*, and *MdMPK6/02G* into pTRV2. The verified plasmids were introduced into *A*. *tumefaciens* strain GV3101 and used to infiltrate ‘Ruixue’ or ‘Fuji’ apples at 165 DAFB. Following five days in dark conditions, the peel surrounding the infiltration location was collected for phenotypic assessment and RT-qPCR with primers listed in Supplemental Table S3.

### *MdWRKY70L* overexpression and knockout in ‘Orin’ calli

*MdWRKY70L* CDS was cloned into pCAMBIA2300 vector for overexpression. CRISPR/Cas9 knockdown targets and corresponding primers (Supplemental Table S3) for *MdWRKY70L* were selected using the website http://crispr.hzau.edu.cn/CRISPR2/. The target single-guide RNA was cloned into pHSE401 and introduced into *A. tumefaciens* LBA4404 cells, which were kept at −80°C until calli transformation.

### Y2H assay

*MdMPK6/02G* CDS was cloned into pGBKT7 and co-transformed with pGADT7 vector into Y2H yeast cells. Simultaneously, *MdWRKY70L* CDS was cloned into the pGADT7 vector. Y2H assays were conducted, as reported previously (Zhang et al., 2023). *MdMPK6/02G* and *MdWRKY70L* interaction was evaluated by observing yeast growth on tryptophan, leucine, histidine, and adenine-deficient medium. Supplemental Table S3 lists all used primers.

### EMSA

*MdWRKY70L* CDS was cloned into pET32a-His and transformed into *E. coli* BL21. The induced proteins were purified and stored at −80°C. The potential MdWRKY70L binding sites in senescence-related genes’ promoter regions were analyzed with PlantCARE software. Biotin-labeled probes, unlabeled competitive probes, and mutant probes were designed for these sites. The binding specificity was confirmed using the LightShift Chemiluminescent EMSA kit (Thermo). Table S3 lists all used primers.

### Dual-LUC reporter analysis

The LUC analysis was executed as reported by Wang et al. (2023). *MdWRKY70L* CDS was inserted into effector vector pGreenII 62-SK driven by CaMV35S promoter. *MdSAG101* and *MdZAT12 p*romoters were cloned into reporter vector pGreenII 0800-LUC. These vectors, along with helper plasmid P19, were introduced into *A. tumefaciens* LBA4404 cells for transient expression in 4-week-old *Nicotiana benthamiana* leaves. The REN sequence in pGreenII 0800-LUC, controlled by the 35S promoter, acted as the positive control. Firefly and Renilla luciferase activities were determined using Infinite M200 (Tecan, Switzerland) with six replicates, and LUC activity 3 days post-infiltration was visualized using an *in vivo* NightOwl II LB983 imaging system (Berthold Technologies, Germany). Table S3 lists all used primers.

### BiFC assay

For the BiFC analysis, *MdMPK6/02G* and *MdWRKY70L* CDS were fused with the N-terminal vector pSPYNE-YFP and C-terminal vector pSPYCE-YFP, respectively. After transformation into *Agrobacterium* cells, they were co-injected into tobacco leaves. The fluorescence signals, indicating protein-protein interaction, were observed under an ultra-high-resolution microscope two days post-injection. Table S3 lists all used primers.

### Firefly LCI assay

For the LCI assay, *MdMPK6/02G* and *MdWRKY70L* CDS were cloned into vector pCAMBIA1300-cLUC and pCAMBIA1300-nLUC, respectively. After transformation into *Agrobacterium* cells, they were co-injected into tobacco leaves. Fluorescence activity was detected in *vivo* using imaging techniques for better clarity. Table S3 lists all used primers.

### Pull-down assay

*MdWRKY70L* and *MdMPK6/02G* CDS were cloned pET-32a (+) and pGEX-4T-1 and transformed into BL21 cells to produce His-tagged and GST-tagged fusion proteins, respectively. These proteins were purified using a commercial protein purification kit (Beyotime Biotechnology, Shanghai, China) and subjected to Western blotting using anti-GST and anti-His antibodies (Abmart, Shanghai, China). Table S3 lists all used primers.

### Protein phosphorylation detection

The assay was conducted in Beijing Bio-Tech Pack Technology Company Ltd. In detail, 10 μg proteins in 100 μL of 50 mmol·L^-^¹ NH₄HCO₃ were reduced with 10 mmol·L^-^¹ DTT for 1 h at 56°C and incubated with 50 mmol·L^-^¹ IAM for 40 minutes in the dark. After that, proteins were digested at 37°C for 4 h with 1% trypsin and 16 h with 2% trypsin. After desalt using a self-packed column, peptides were dried at 45°C using a vacuum centrifuge and re-solubilized in 0.1% formic acid. After centrifugation at 13,200 rpm for 10 min at 4°C, samples were subjected to mass spectrometry analysis for over 66 min using 100 μm i.d. × 180 mm pre-packed 3 μm Reprosil-Pur 120 C18-AQ column with 0.1% formic acid as mobile phase A and 0.1% formic acid in 80% ACN as mobile phase B at flow rate of 600 nL·min^-^¹.

### Validation of protein phosphorylation *in vitro*

*MdWRKY70L* phosphorylation at Ser199 was identified through IP/MS analysis. For further *in vitro* validation, the site was mutated to aspartic acid. WT and mutated MdWRKY70L and CAMdMPK6/02G were cloned into pGEX4T-1-GST, expressed in *E. coli* BL21 cells, and purified, respectively. The purified Wt and mutated MdWRKY70L proteins were mixed with CAMdMPK6/02G, respectively, at 1:5 ratio and incubated in kinase reaction buffer at 25°C for 30 min. MdWRKY70L phosphorylation by *MdMPK6/02G* was assessed through Western blotting. Table S3 lists all used primers.

### Western blotting

Western blotting was executed previously described (Wang et al., 2023) using anti-GFP, anti-GST, and anti-phos antibodies from Abmart Medical Technology (Shanghai, China) Co., Ltd.

### Statistics

All experiments were executed with three biological repeats. Data were processed using Microsoft Excel 2010, SigmaPlot 13, and Origin 2017 and compared using one-way analysis of variance (ANOVA) and student’s *t* test using SPSS 24.0 with *P* < 0.05 considered significant.

## ACCESSION NUMBERS

All RNA-seq data was submitted to https://www.ncbi.nlm.nih.gov/sar under accession number PRJNA861871. Other sequencing data can be downloaded from http://plants.ensembl.org/index.html under MD09G0105800 for *MdWRKY1*, MD13G0059600 for *MdWRKY3*, MD03G0162000 for *MdWRKY31*, MD03G0048200 for *MdWRKY24*, MD13G0134000 for *MdWRKY48*, MD05G0248800 for *MdWRKY65*, MD09G0202900 for *MdWRKY69*, MD01G0136400 for *MdWRKY70L*, MD13G0068300 for *MdWRKY72A*, MD13G0108800 for *MdWRKY75*, MD15G0034900 for *MdWRKY76*, MD09G0034000 for *MdSAG101*, MD14G0164000 for *MdEDS1*, MD12G0174000 for *MdCBP60F*, MD13G0103200 for *MdCYP76B6*, MD17G0093500 for *MdACO1*, MD14G0097100 for *MdACS1*, MD11G0144200 for *MdAAO1*, MD04G0166700 for *MdLOX1.5*, and MD07G0159300 for *MdZAT12*.

## SUPPLEMENTARY DATA

**Supplemental Figure S1.** Determination of antioxidant capacity and ROS enzyme activity in different parts of apple fruits during senescence. (a) Chlorophyll content. (b) Total polyphenols. (c) Total flavonoids. (d) Total flavanols. (e) Total antioxidant activity. (f) SOD activity. (g) POD activity. (h) CAT activity. Data shown are mean ± standard error with different letters denoting *P* < 0.05 (Student’s *t* test).

**Supplemental Figure S2.** Senescence-related gene expression levels in fruits at various stages. Different letters represent significant differences at *P* < 0.05 (Student’s *t* test).

**Supplemental Figure S3.**Senescence-related gene expression levels after *MdWRKY70L* transfection into apple and calli. (a) Senescence-related gene expression levels after instant insertion of *MdWRKY70L* into apple. (b) Senescence-related gene expression levels after stable insertion of *MdWRKY70L* into calli. Data shown are mean ± standard error with different letters denoting *P* < 0.05 (Student’s *t* test).

**Supplemental Figure S4.** ROS biosynthesis in stable transgenic calli. (a) Calli phenotype after NBT staining for O₂⁻· with darker colors representing higher content) and DAB for H₂O₂ with browner colors representing higher content. (b) O₂⁻· content. (c) H₂O₂ content. Data shown are mean ± standard error with different letters denoting *P* < 0.05 (Student’s *t* test).

**Supplemental Figure S5.** Protein posttranslational modification detection using liquid chromatography-tandem mass spectrometry. (a) Total ion flow chromatogram of wild-type calli. (b) Total ion flow chromatogram of stable transgenic *MdWRKY70L* calli.

**Supplemental Figure S6.** Verification of protein phosphorylation stability in *vivo*.

**Supplemental Figure S7.**MdWRKY70L phosphorylation at Ser199 by MdMPK6/02G accelerated fruit senescence. Scale bar = 2 cm. Apple images were digitally processed for comparison.

**Supplemental Table S1.** Identification of phosphorylated peptides.

**Supplemental Table S2.** RT-qPCR primers.

**Supplemental Table S3.** Primers for transgene construction, CRISPR/Cas9-based knockout, electrophoretic mobility shift assay, luciferase assay, yeast two-hybrid assay, bimolecular fluorescence complementation assay, and luciferase complementation imaging.

## ACKNOWLEDGMENTS

We extend our gratitude to Professor Xuesen Chen’s team at Shandong Agricultural University for providing essential carriers and experimental materials. We also thank Guangzhou Genedenovo Biotechnology Co., Ltd. for their support with sequencing and bioinformatics analysis. Special thanks to Topedit (https://www.topeditsci.com/) for English polishing.

## AUTHOR CONTRIBUTIONS

Z.Z. and H.W. conceived the study. H.W. and S.Z. executed the experiments, provided reagents and materials, and analyzed the data. H.W. S.Z. and Z.Z. prepared the paper.

## REFERENCES

Buchanan-Wollaston V, Page T, Harrison E, Breeze E, Lim P, Nam H, Lin J, Wu S, Swidzinski J, Ishizaki K, Leaver C. Comparative transcriptome analysis reveals significant differences in gene expression and signalling pathways between developmental and dark/ starvation-induced senescence in *Arabidopsis*. Plant Journal. 2005:42(4):567–585. 10.1111/j.1365-313X.2005.02399.x

Chen L, Xiang S, Chen Y, Li D, Yu D. *Arabidopsis* WRKY45 interacts with the DELLA protein RGL1 to positively regulate age-triggered leaf senescence. Molecular Plant. 2017:10(9):1174–1189. 10.1016/j.molp.2017.07.008

Chen Q, Yan J, Tong T, Zhao P, Wang S, Zhou N, Cui X, Dai M, Jiang Y, Yang B. A NAC087 transcription factor positively regulates age-dependent leaf senescence through modulating the expression of multiple target genes in *Arabidopsis*. Journal of Integrative Plant Biology. 2023:65(4):967–984. 10.1111/jipb.13434

Decros G, Baldet P, Beauvoit B, Stevens R, Flandin A, Colombié S, Gibon Y, Pétriacq P. Get the balance right: ROS homeostasis and redox signalling in fruit. Frontiers in Plant Science. 2019:10(9):1091. 10.3389/fpls.2019.01091

Giovannoni J. Molecular biology of fruit maturation and ripening. Annual Review of Plant Biology. 2001:52(1):725–749. 10.1146/annurev.arplant.52.1.725

Guan Y, Meng X, Khanna R, LaMontagne E, Liu Y, Zhang S. Phosphorylation of a WRKY transcription factor by MAPKs is required for pollen development and function in *Arabidopsis*. Plos Genetics. 2014:10(5):e1004384. 10.1371/journal.pgen.1004384

Guo P, Li Z, Huang P, Li B, Fang S, Chu J, Guo H. A tripartite amplification loop involving the transcription factor WRKY75, salicylic acid, and reactive oxygen species accelerates leaf senescence. The Plant Cell. 2017:29(11):2854–2870. 10.1105/tpc.17.00438

Han M, Kim C, Lee J, Lee S, Jeon J. OsWRKY42 represses OsMT1d and induces reactive oxygen species and leaf senescence in rice. Molecules and Cells. 2014:37(7):532–539. 10.14348/molcells.2014.0128

Hu L, Ye M, Li R, Zhang T, Zhou G, Wang Q, Lu J, Lou Y. The rice transcription factor WRKY53 suppresses herbivore-induced defenses by acting as a negative feedback modulator of mitogen-activated protein kinase activity. Plant Physiology. 2015:169(4):2907–2921. 10.1104/pp.15.01090

Huang W, Hu N, Xiao Z, Qiu Y, Yang Y, Yang J, Mao X, Wang Y, Li Z, Guo H. A molecular framework of ethylene-mediated fruit growth and ripening processes in tomato. The Plant Cell. 2022:34(9):3280–3300. 10.1093/plcell/koac146

Jiang G, Yan H, Wu F, Zhang D, Zeng W, Qu H, Chen F, Tan L, Duan X, Jiang Y. Litchi fruit LcNAC1 is a target of LcMYC2 and regulator of fruit senescence through its interaction with LcWRKY1. Plant Cell Physiology. 2017:58(6):1075– 1089. 10.1093/pcp/pcx054

Kuang J, Chen J, Luo M, Wu K, Sun W, Jiang Y, Lu W. Histone deacetylase HD2 interacts with ERF1 and is involved in longan fruit senescence. Journal of Experimental Botany. 2012:63(1):441–454. 10.1093/jxb/err290

Li H, Ding Y, Shi Y, Zhang X, Zhang S, Gong Z, Yang S. MPK3- and MPK6- mediated ICE1 phosphorylation negatively regulates ICE1 stability and freezing tolerance in *Arabidopsis*. Developmental Cell. 2017:43(5):630–642. 10.1016/j.devcel.2017.09.025

Li X, Guo W, Li J, Yue P, Yue P, Bu H, Jiang J, Liu W, Xu Y, Yuan H, Li T, Wang A. Histone acetylation at the promoter for the transcription factor PuWRKY31 affects sucrose accumulation in pear fruit. Plant Physiology. 2020:182(4):2035– 2046. 10.1104/pp.20.00002

Lokdarshi A, Guan J, Camacho R, Cho S, Morgan P, Leonard M, Shimono M, Day B, Arnim A. Light activates the translational regulatory kinase GCN2 via reactive oxygen species emanating from the chloroplast. The Plant Cell. 2020:32(4):1161–1178. 10.1105/tpc.19.00751

Mao G, Meng X, Liu Y, Zheng Z, Chen Z, Zhang S. Phosphorylation of a WRKY transcription factor by two pathogen-responsive MAPKs drives phytoalexin biosynthesis in *Arabidopsis*. The Plant Cell. 2011:23(4):1639–1653. 10.1105/tpc.111.084996

Miao Y, Laun T, Zimmermann P, Zentgraf U. Targets of the WRKY53 transcription factor and its role during leaf senescence in *Arabidopsis*. Plant Molecular Biology. 2004:55(6):853–867. 10.1007/s11103-005-2142-1

Miao Y, Zentgraf U. The antagonist function of Arabidopsis WRKY53 and ESR/ESP in leaf senescence is modulated by the jasmonic and salicylic acid equilibrium. The Plant Cell. 2007:19(3):819–830. 10.1105/tpc.106.042705

Mittler R. ROS are good. Trends in Plant Science. 2017:22(1):11–19. 10.1016/j.tplants.2016.08.002

Mittler R, Zandalinas S, Fichman Y, Breusegemrank F. Reactive oxygen species signalling in plant stress responses. Nature Reviews Molecular Cell Biology. 2022:23(10):663–679. 10.1038/s41580-022-00499-2

Muñoz P, Munné-Bosch S. Photo-oxidative stress during leaf, flower and fruit development. Plant Physiology. 2018:176(2):1004–1014. 10.1104/pp.17.01127

Shan W, Kuang J, Chen L, Xie H, Peng H, Xiao Y, Li X, Chen W, He Q, Chen J, Lu W. Molecular characterization of banana NAC transcription factors and their interactions with ethylene signaling component EIL during fruit ripening. Journal of Experimental Botany. 2012:63(14):5171–5187. 10.1093/jxb/ers178

Shinozaki Y, Nicolas P, Fernandez-Pozo N, Ma Q, Evanich D, Shi Y, Xu Y, Zheng Y, Snyder S, Martin L, Ruiz-May E, Thannhauser T, Chen K, Domozych D, Catalá C, Fei Z, Mueller L, Giovannoni J, Rose J. High-resolution spatiotemporal transcriptome mapping of tomato fruit development and ripening. Nature Communications. 2018:9(1):364. 10.1038/s41467-017-02782-9

Sun T, Zhang Y. MAPK kinase cascades in plant development and immune signaling. EMBO Reports. 2021:23(2):e53817. 10.15252/embr.202153817

Tian S, Qin G, Li B. Reactive oxygen species involved in regulating fruit senescence and fungal pathogenicity. Plant Molecular Biology. 2013:82(6):593–602. 10.1007/s11103-013-0035-2

Tian X, Li X, Zhou W, Ren Y, Wang Z, Liu Z, Tang J, Tong H, Fang J, Bu Q. Transcription factor OsWRKY53 positively regulates brassinosteroid signaling and plant architecture. Plant Physiology. 2017:175(3):1337–1349. 10.1104/pp.17.00946

Wang H, Zhang S, Fu Q, Wang Z, Liu X, Sun L, Zhao Z. Transcriptomic and metabolomic analysis reveals a protein module involved in pre-harvest apple peel browning. Plant Physiology. 2023:192(3):2102–2122. 10.1093/plphys/kiad064

Wang P, Liu W, Han C, Wang S, Bai M, Song C. Reactive oxygen species: Multidimensional regulators of plant adaptation to abiotic stress and development. Journal of Integrative Plant Biology. 2024:66(3):1–38. 10.1111/jipb.13601

Wang Y, Tian S, Xu Y. Effects of high oxygen concentration on pro-and anti-oxidant enzymes in peach fruits during postharvest periods. Food Chemistry. 2004:91(6):99–104. https://api.semanticscholar.org/CorpusID:85092154

Xiao Y, Chen J, Kuang J, Shan W, Xie H, Jiang Y, Lu W. Banana ethylene response factors are involved in fruit ripening through their interactions with ethylene biosynthesis genes. Journal of Experimental Botany. 2013:64(8):2499–2510. 10.1093/jxb/ert108

Yang L, Ye C, Zhao Y, Cheng X, Wang Y, Jiang Y, Yang B. An oilseed rape WRKY-type transcription factor regulates ROS accumulation and leaf senescence in *Nicotiana benthamiana* and *Arabidopsis* through modulating transcription of RbohD and RbohF. Planta. 2018:247(6):1323–1338. 10.1007/s00425-018-2868-z

Zhang L, Wang L, Fang Y, Gao Y, Yang S, Su J, Ni J, Teng Y, Bai S. Phosphorylated transcription factor PuHB40 mediates ROS-dependent anthocyanin biosynthesis in pear exposed to high light. The Plant Cell. 2024:36(9):3562–3583. 10.1093/plcell/koae167

Zhang Y, Liu Z, Wang X, Wang J, Fan K, Li Z, Lin W. DELLA proteins negatively regulate dark-induced senescence and chlorophyll degradation in *Arabidopsis* through interaction with the transcription factor WRKY6. Plant Cell Reports. 2018:37(7):981–992. 10.1007/s00299-018-2282-9

Zhang Y, Wang Y, Wei H, Li N, Tian W, Chong K, Wang L. Circadian evening complex represses jasmonate-induced leaf senescence in *Arabidopsis*. Molecular Plant. 2018:11(2):326–337. 10.1016/j.molp.2017.12.017

Zhao J, Quan P, Liu H, Li L, Qi S, Zhang M, Zhang B, Li H, Zhao Y, Ma B, Han M, Zhang H, Xing L. Transcriptomic and metabolic analyses provide new insights into the apple fruit quality decline during long-term cold storage. Journal of Agricultural and Food Chemistry. 2020:68(16):4699–4716. 10.1021/acs.jafc.9b07107

Zhao M, Wang J, Shan W, Fan J, Kuang J, Wu K, Li X, Chen W, He F, Chen J, Lu W. Induction of jasmonate signaling regulators MaMYC2s and their physical interactions with MaICE1 in methyl jasmonate-induced chilling tolerance in banana fruit. Plant Cell Environment. 2013:36(1):30–51. 10.1111/j.1365-3040.2012.02551.x

Zhu G, Wan S, Huang Z, Zhang S, Liao Q, Zhang C, Lin T, Qin M, Peng M, Yang C, Cao X, Han X, Wang X, Knaap E, Zhang Z, Cui X, Klee H, Fernie A, Luo J, Huang S. Rewiring of the fruit metabolome in tomato breeding. Cell. 2018:172(1-2):249–261. 10.1016/j.cell.2017.12.019

Zhu L, Chen L, Wu C, Shan W, Cai D, Lin Z, Wei W, Chen J, Lu W, Kuang J. Methionine oxidation and reduction of the ethylene signaling component MaEIL9 are involved in banana fruit ripening. Journal of Integrative Plant Biology. 2023:65(1):150–166. 10.1111/jipb.13363

